# TEINet: a deep learning framework for prediction of TCR-epitope binding specificity

**DOI:** 10.1101/2022.10.20.513029

**Authors:** Yuepeng Jiang, Miaozhe Huo, Shuai Cheng Li

## Abstract

The adaptive immune response to foreign antigens is initiated by T-cell receptor (TCR) recognition on the antigens. Recent experimental advances have enabled the generation of a large amount of TCR data and their cognate antigenic targets, allowing machine learning models to predict the binding specificity of TCRs. In this work, we present TEINet, a deep learning framework that utilizes transfer learning to address this prediction problem. TEINet employs two separately trained encoders to transform TCR and epitope sequences into numerical vectors, which are subsequently fed into a fully connected neural network to predict their binding specificities. A major challenge for binding specificity prediction is the lack of a unified approach to sample negative data. Here, we first assess the current negative sampling approaches comprehensively and suggest that the *Unified Epitope* is the most suitable one. Subsequently, we compare TEINet with three baseline methods and observe that TEINet achieves an AUROC of 0.760, which outperforms baseline methods by 6.4-26%. Furthermore, we investigate the impacts of the pretraining step and notice that excessive pretraining can adversely affect model performance. Our results and analysis show that TEINet can make an accurate prediction using only the TCR sequence (CDR3*β*) and the epitope sequence, providing novel insights to understand the interactions between TCRs and epitopes. TEINet is available at https://github.com/jiangdada1221/TEINet.

## Introduction

T cells are critical for the adaptive immune system, providing protection against a wide range of pathogens. To recruit T cells in an immune response, the T cell receptors (TCRs) on their surface have to recognize a non-self-immunogenic peptide (epitope) presented in the context of major histocompatibility complex molecules (MHC). The generation of these protein receptors arises mainly from the quasirandom somatic V(D)J recombination process which theoretically can produce extremely high TCR diversity of 10^15^-10^20^ in an individual, each with unique recognition capacity for antigens [1]. Understanding the mechanisms that govern the interaction between TCR and peptide-MHC (pMHC) is considered an essential step toward personalized immunotherapy and the development of targeted vaccines.

Recent advancements in the high-throughput tetramer-associated T cell receptor sequencing technique [2] and other experimental approaches such as tetramer analysis [3] and T-scan [4] have enabled the generation of an increasing amount of data recording the binding of TCR and epitope. More and more interaction pairs are consistently being generated and stored in publicly available databases such as VDJdb [5], IEDB [6] and McPAS-TCR [7]. However, the available data are still scant compared to the theoretical TCR diversity. Further, the TCR-epitope paired data are imbalanced, as a single epitope is often linked by many TCRs. Both of them pose challenges to the development of *in silico* predictive methods.

Machine learning-based methods are able to capture the potential laws of TCR-epitope binding from a large amount of experimental data. With the help of advanced machine learning models, several computational methods have been proposed to assess the binding of a TCR and a pMHC (epitope). Previously, a branch of research focused on designing epitope-specific models with the aim of learning the pattern of TCRs binding to the same epitope. These models range from simple sequence alignment-based methods [8] to more complex machine learning models including random forest (e.g. TCRex [9]) and the Gaussian process classifier TCRGP [10]. However, they all share two downsides: each epitope needs a specific model trained separately; each model requires abundant training samples of epitope-specific TCRs, which are not always readily available.

To fulfill the need to predict the binding specificity of any TCR-epitope pair, previous studies have proposed generic models, which exploit the two-tower architecture to encode both the TCRs (CDR3*β*) and the pMHCs (epitope) [11–17]. These generic models can fully capitalize on the currently available paired data to unlock the binding patterns between TCRs and epitopes, and transfer the knowledge learned from paired samples of epitopes with sufficient binding TCRs to those with sparse linking TCRs. Current models have shown moderate predictive performance and demonstrated promising potential in understanding cancer progression, prognosis, and responsiveness to immunotherapy. For example, Dash *et al*. developed TCRdist [17] based on sequence similarity weighted distances; Moris *et al*. proposed a CNN-based model ImRex [11]; Weber *et al*. introduced TITAN that encodes epitopes at the atomic level with SMILES sequences using a pretrained deep learning model. Furthermore, Lu *et al*. presented pMTNet [14] that encodes TCRs and pMHCs by two respective pre-trained deep learning models and applied pMTNet to investigate tumor progression and response to immunotherapy treatment. In particular, transfer learning is becoming a prior technique to develop advanced deep learning models for binding prediction since it helps leverage the knowledge from other pretraining tasks with abundant data and transfer it to the binding prediction task. For instance, TITAN, NetTCR [15], and pMTnet utilize pre-trained encoders. However, the impact of the pretraining step on the final performance of predicting TCR specificity remains undiscovered.

To train and evaluate supervised models, both positive and negative samples (TCRs and epitopes that do not interact with each other) are required. However, the public TCR-epitope interaction datasets only collect positive samples, which potentially poses a challenge in model training and evaluation. The method of generating negative samples based on the existing TCR and epitope pairs directly affects the model performance. Currently, there are four major strategies for generating negative samples: (1) *Reference TCR* [15, 18]; (2) *Random TCR* [14]; (3) *Random Epitope* [12, 16, 19]; (4) *Unified Epitope* [11, 20]. Different models might adopt different negative sampling strategies for this task, which make it difficult to fairly compare their performance. More importantly, which strategy leads to a better generalized model has not been explored and remains an open question.

In this work, we present TEINet for the prediction of the specificity of TCR binding, using the CDR3*β* chain of TCR and the epitope sequence within the pMHC complex. Following the concept of transfer learning, TEINet employs two separate pretrained encoders to convert TCRs and epitopes into numerical vectors, utilizing the architecture of recurrent neural networks to handle a variety of sequence lengths. We first contrast the four negative sampling strategies applied in the previous work to select the superior one. Next, we systematically validated TEINet using a large-scale TCR-epitope paired dataset and two independent validation datasets. The results demonstrated the enhancement in accuracy made over previous work. We also investigated the impact of the pretraining step on the final binding specificity prediction task. Overall, TEINet serves as a reliable computational tool for addressing the long-standing problem of predicting the TCR-epitope interaction.

## Methods

### Dataset

The CDR3 regions of TCR*β* chains are located in the center of the paratope and are considered as the key determinant of specificity in antigen recognition [21]. Although CDR3-*α* and -*β* synergistically drive TCR-epitope recognition [17, 22, 23], the current available databases still record mostly *β*chain paired samples. Thus, we restrict ourselves to CDR3*β* chain sequences in this study. Besides, with the aim of developing a general model that is suitable for most cases, we took the epitope sequence inside the pMHC complex as its representation. In order to construct a large and diverse dataset, we combined the data recorded in VDJdb database [5], McPAS database [7], and the data collected by Lu *et al*. [14] together.

The data from VDJdb was downloaded from its public website (https://vdjdb.cdr3.net/) on April 5, 2022. It consists of 89,321 curated pairs of CDR3 *α*/*β* sequences along with their binding epitopes and MHC classes, covering three species. We selected only human TCR sequences, removed duplicate cases, restricted only MHC class I entries, and only kept the CDR3*β* and epitope sequences whose lengths lie between 5-30 and 7-15 amino acids, respectively. After all these filterings, this dataset was reduced to 35,560 unique CDR3*β*-epitope pairs, among which 33,258 TCRs are assigned to 159 epitopes.

The McPAS-TCR dataset [7] originally contains 39,664 pairs (http://friedmanlab.weizmann.c.il/McPAS-TCR/) and Lu *et al*. collected a total of 32,607 paring data from a series of previous publications and four chromium single-cell immune profiling solution datasets. We performed the same preprocessing step on these two datasets and removed all TCR sequences with ambiguous amino acids (B, J, O, U, X). Then, these three datasets were merged together, followed by two additional filtering steps: removal of duplicate pairs and exclusion of epitopes with less than 10 associated TCR sequences since this merged dataset is highly imbalanced. At last, we constructed a large dataset with 44,682 pairs of TCRs and epitopes, among which 41,610 TCRs are linked to 180 epitopes. An overview of the dataset is shown in Fig. 1.

**Figure 1:**
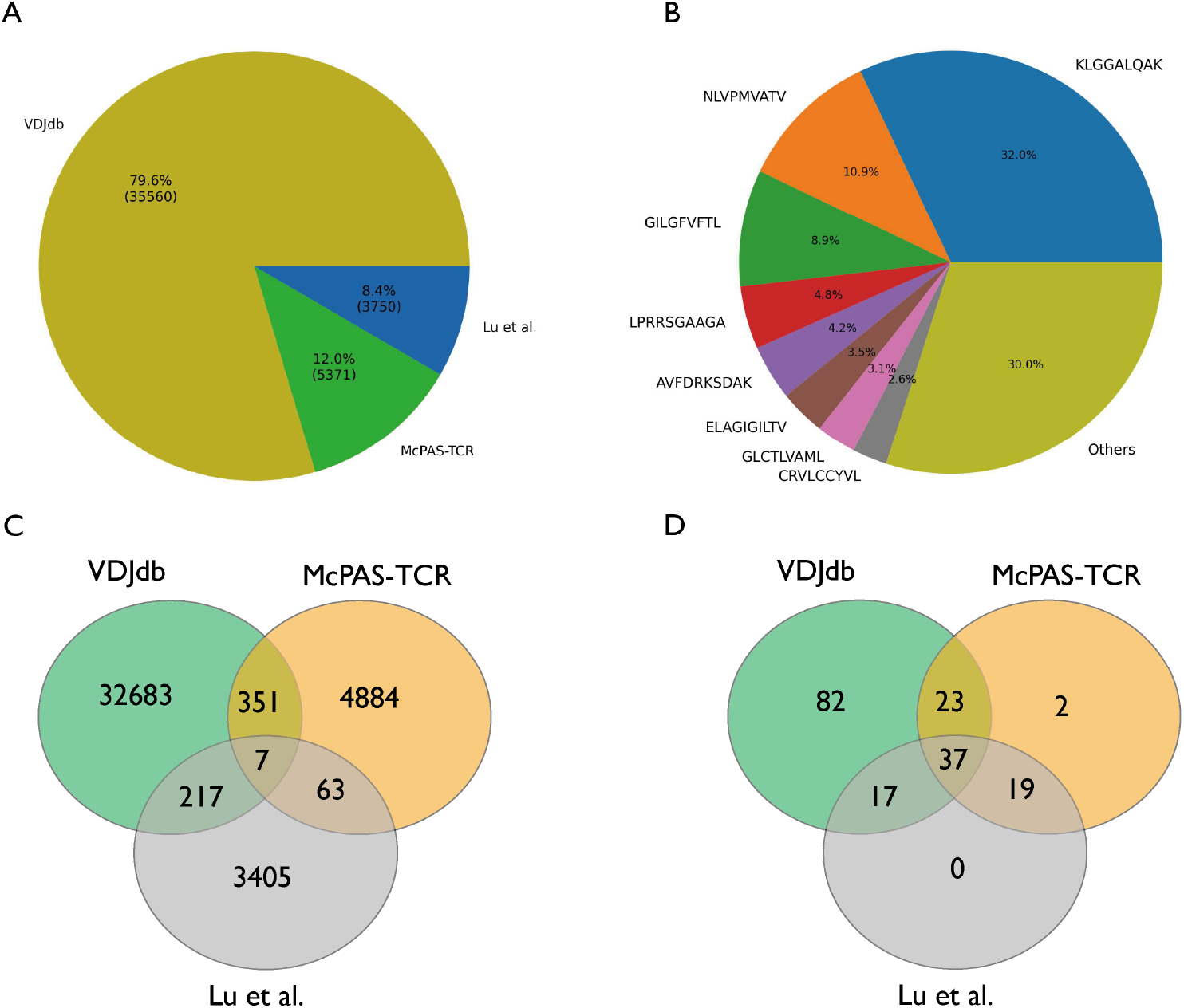
Overview of our constructed dataset. (A) The source of the paired samples in the dataset. (B) The number of the epitope-associated TCRs. Most TCRs are linked to a small group of epitopes. (C and D) Venn diagrams showing the number of (C) TCRs and (D) epitopes contributed by each source dataset.

### Negative sampling strategies

Since the TCR-epitope dataset contains only positive samples, in order to train a generalized and robust supervised model, the negative samples are required and should be generated via a biologically and computationally plausible manner to serve as an unbiased estimate of the actual distribution of non-binding pairs. For a positive sample 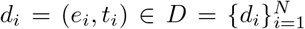, where *e*_*i*_ and *t*_*i*_ are the interacting epitope and TCR for sample *i*, the corresponding negative samples are generated through four major sampling strategies (Fig. 2A):

**Figure 2:**
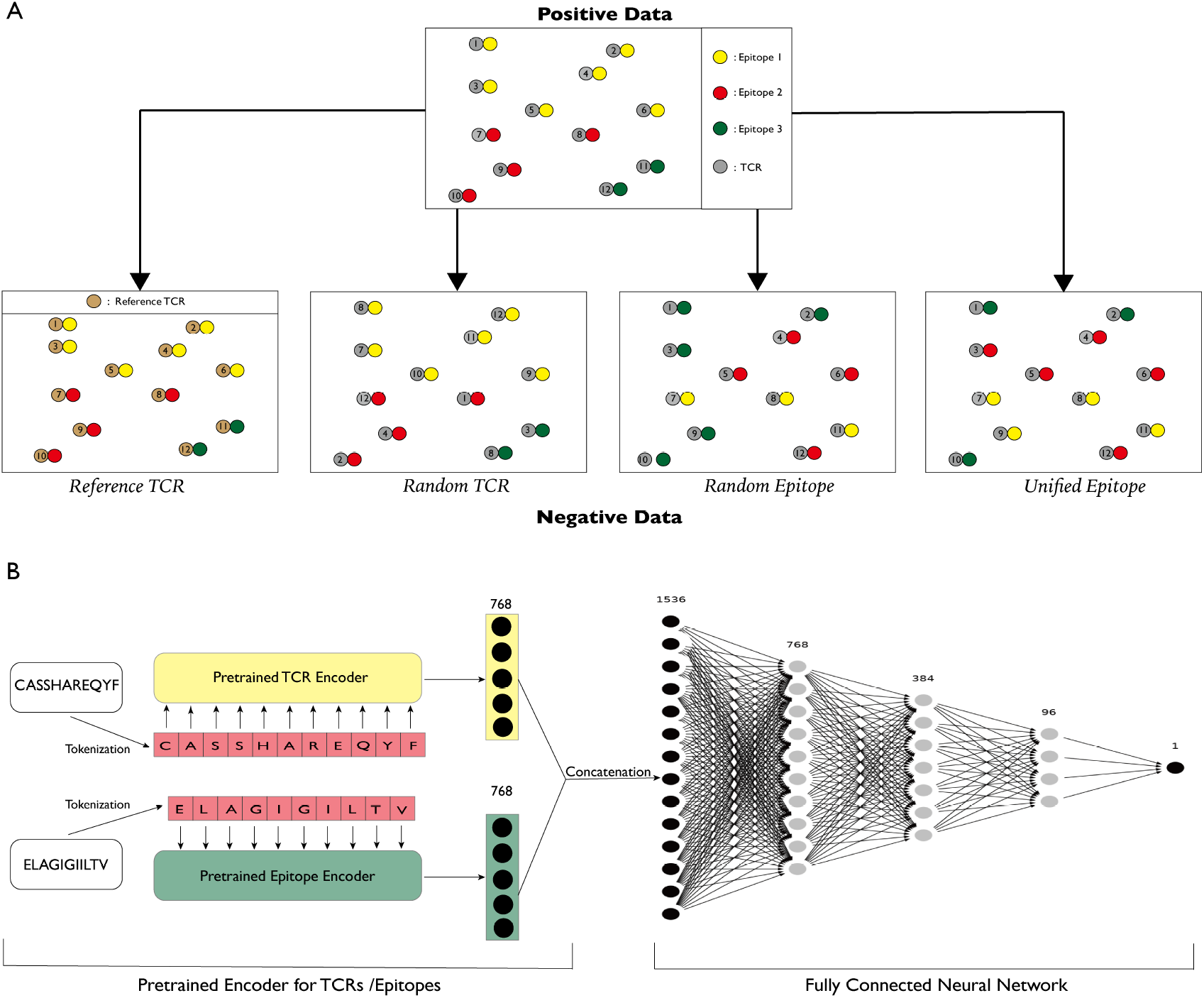
Illustration of each negative sampling strategy and the overall workflow of TEINet. (A) Sketch map of the four negative sampling strategies. In this example, there are in total 12 TCR-epitope binding pairs, with 3 different epitopes (depicted in yellow, red, and green) linking to 6, 4, and 2 TCRs, respectively. For *Reference TCR* strategy, the TCRs are randomly sampled from a reference TCR dataset inside which TCRs are considered unable to bind epitopes in the positive data. Here, we choose to generate the same number of pairs in negative data for demonstration. (B) General workflow of TEINet. TEINet is a two-stage deep learning model using transfer learning. At the first pretraining stage, two TCRpeg models are trained separately to learn the sequence pattern of TCRs and epitopes, and produce numerical encodings for them when the pretraining process is completed. At the next stage, encodings of TCRs and epitopes are concatenated together and output into a fully connected neural network to leverage the information from each part and make predictions accordingly.

- *Reference TCR*. In this setting, *e*_*i*_ is combined with TCRs that are sampled uniformly from the reference TCR dataset *R* = {*t*_*j*_}. The negative samples for *e*_*i*_ are then represented as 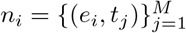, where *t*_*j*_ ∈ *R* and M is the number of negatives samples for a given positive sample [15, 18]. This approach stands upon the assumption that TCRs from the reference dataset are unlikely to bind epitopes in the positive dataset. The reference TCRs were obtained from Montemurro *et al*. [15] where these TCRs had been exposed to all tested pMHC multimers and no binding signals were detected.
- *Random TCR*. For this sampling approach, the negative TCRs for *e*_*i*_ are sampled uniformly from the set of TCRs in the positive binding pairs while excluding its known true TCR binding partner(s) [14]. The negative samples are then represented as 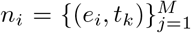, where *t*_*k*_ ∈ {*t*_*i*_} and (*e*_*i*_, *t*_*k*_) ∉ *D*.
- *Random Epitope*. In this strategy, *t*_*i*_ is combined with epitopes sampled uniformly from all epitopes without its true epitope binder(s) [12, 16, 19]. The sampled negative pairs are 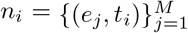 with *e*_*j*_ ∈ {*e*_*i*_} and (*e*_*j*_, *t*_*i*_) ∉ *D*.
- *Unified Epitope*. Compared to *Random Epitope*, the only difference of *Unified Epitope* is that, the epitopes are sampled according to their frequency distributions in the positive dataset [11, 20]; i.e. 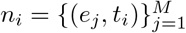 and *P*_*pos*_(*e*_*j*_) ∼ *P*_*neg*_(*e*_*j*_). This strategy ensures that the frequencies of epitopes are unified in the negative data and positive data.

A systematical comparison between these four negative sampling strategies is an urgent need for benchmarking different models and guiding the development of accurate and generalized models in future works. To address this demand, we referred to the field of recommender system and selected three evaluation metrics that can be calculated without the attendance of negative samples.

#### Precision@k and Recall@k

These two metrics measure the exactness and completeness of the top k binding predictions for a given TCR. Assume that a TCR *t*_*i*_ in the test set 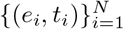 can bind to a number of *b*_*i*_ epitopes (due to cross-reactivity), and a number of *m*_*i*_ true interacting pairs 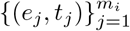lie in the top k predictions, then these two metrics are defined as follows:

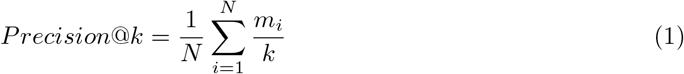

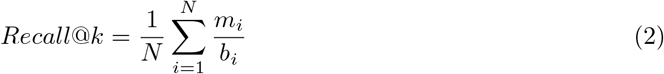

where *N* is the total number of TCRs in the test set. A higher value of *Precision@k* indicates that the more true binding pairs can be found among the top k predicted pairs; And a higher value of *Recall@k* suggests a higher proportion of predicted binding pairs over all the true binding pairs.

#### NDCG@k

The previous two metrics overlook the order of the predictions since the ranking of the true predicted binding pairs does not affect the values of both metrics as long as they are in the top k predictions. The Normalized Discounted Cumulative Gain (NDCG) measures how relevant the predictions are and how good the ordering is, which is calculated by:

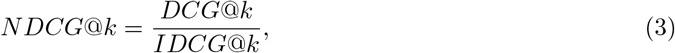

where the definitions and formulas of DCG@k and IDCG@k are described in Supplementary Text S1. Overall, these three metrics are complementary to each other and help to determine the superiority of the four negative sampling strategies. A higher value of any of the three metrics indicates a better model performance.

### Pretrained encoder

To numerically encode TCR and epitopes, we capitalized on the transfer learning technique. We have earlier proposed an autoencoder model, TCRpeg [24] that utilizes a recurrent neural network with GRU layers to characterize the TCR repertoires and demonstrated that it can produce high-quality vector encodings for TCR sequences. As an autoencoder model, TCRpeg is capable of capturing key features of sequence input via unsupervised learning of mapping between the latent space and sequence space, and more importantly, using TCRpeg for pretraining only needs plain amino acid sequences which are currently abundant. In addition, unlike the encoders in TITAN or pMTNet, TCRpeg can process sequences of arbitrary lengths without the need to pad them to a fixed length. Thus, we decided to employ two separate pretrained TCRpeg models as the encoders for TCRs and epitopes, respectively. A detailed description of TCRpeg is given in Supplementary Text S2.

To pretrain TCRpeg for encoding TCRs (TCRpeg-TCR), we fed TCRpeg with 10^6^ TCR sequences collected from Emerson *et al*. [25]. We set the feature size of TCRpeg to 768 and trained it for 20 epochs by minimizing the cross-entropy loss between the output soft-maxed logits and the one-hot encoded representation of the input sequences. For encodings of epitopes, we trained another TCRpeg model (TCRpeg-Epi) with the identical architecture of TCRpeg-TCR using 362,456 unique epitope sequences collected from Mei *et al*. [26] with lengths ranging from 8 to 14 amino acids. Details on the pretraining process of TCRpeg are elaborated in Supplementary Text S3.

### Model architecture

Figure 2B delineates an overview of the architecture of TEINet. Conceptually, the complex task of predicting the TCR-epitope interaction is decomposed into two steps to lower the difficulty level of the final prediction task. First, two encoding networks are pretrained so that the amino acid sequences of TCRs and epitopes could be represented by numerical vectors. Next, we concatenated these two vector encodings to form the final representations for TCR-epitope pairs. In the final step, we built a fully connected neural network (FCN) on top of these combined vector encodings to fuse the knowledge extracted from TCRs and epitopes. Specifically, the FCN consists of three hidden layers with 768, 384, and 96 neurons with the dropout [27] rate set to 0.15 to prevent overfitting. Before feature concatenation, we employed the layer normalization [28] to numerically stabilize each group of features. All neurons use the scaled exponential linear unit (SELU [29]) activation function, except for the output neuron which applies the sigmoid activation function.

### Model training

TEINet was implemented in Python 3.6 and built on the deep learning framework PyTorch [30]. TEINet was trained and evaluated under a 5-fold cross-validation procedure. Instead of inferring TEINet on a static dataset with negative pairs sampled prior to the training process, we adopted a dynamic sampling strategy: the negative examples are sampled on the fly at each training step using the sampling strategies described in the previous section. This dynamic sampling strategy demonstrates improved performance over static training (Supplementary Figure S1). For all experiments in this work, the negative pairs were sampled 10 times more than positive pairs. TEINet optimized binary cross entropy loss with Adam algorithm [31] and an initial learning rate of 1 × 10^−3^. The model was trained for 30 epochs with a batch size of 48. The learning rate was reduced at the 21st and 27th epoch by a factor of 0.1.

## Results

### Comparison of different negative sampling strategies

We trained TEINet with each negative sampling method and observed that they could achieve performance in different scales (Supplementary Figure S2). For example, using *Reference TCR* leads to an average AUROC (area under the receiver operating characteristic) of 0.797, whereas the performance achieves an AUROC of 0.934 under *Random Epitope*, which is unexpectedly high yet useless.

We first compared the three negative sampling methods: *Random TCR, Reference TCR*, and *Unified Epitope*. The negative data generated by these methods possess similar frequency distributions of epitopes with those in the positive data. Table 1 shows the Precision, Recall, and NDCG of each schema using the TEINet. These results first demonstrated that *Random TCR* and *Reference TCR* obtained similar performance, indicating that sampling TCRs from the reference TCR pool or TCRs in positive data have a comparable effect on the model training. *Reference TCR* is slightly better than *Random TCR*, as TCRs drawn from another sequence pool constructed from healthy donors are less likely to interact with epitopes than shuffled TCRs from the positive data; i.e., *Random TCR* might produce more false negative pairs. *Unified Epitope* achieved superior performance among these three strategies by a large margin. It indicates that *Unified Epitope* can help develop a more robust and generalized model for the TCR-epitope interaction prediction task. We attributed its superior performance to the uniformity of the distribution of TCRs and epitopes across positive and negative data.

**Table 1:** The Precision, Recall, and NDCG of each negative sampling method.

| Method | Precision@3 | Recall@3 | NDCG@3 | Precision@10 | Recall@10 | NDCG@10 |
| --- | --- | --- | --- | --- | --- | --- |
| *Reference TCR* | 0.093±0.002 | 0.275±0.007 | 0.255±0.006 | 0.036±0.001 | 0.356±0.009 | 0.284±0.007 |
| *Random TCR* | 0.085±0.002 | 0.251±0.004 | 0.226±0.004 | 0.033±0.001 | 0.322±0.003 | 0.252±0.003 |
| *Unified Epitope* | 0.129±0.004 | 0.380±0.012 | 0.334±0.011 | 0.052±0.001 | 0.506±0.014 | 0.380±0.012 |
| *Random Epitope* | 0.192±0.002 | 0.567±0.006 | 0.484±0.006 | 0.081±0.001 | 0.788±0.002 | 0.565±0.005 |

We next contrasted *Unified Epitope* with *Random Epitope*. It seems that *Random Epitope* is a perfect sampling strategy since it achieved extremely high values of Precision, Recall, and NDCG (Table 1), and achieved an average AUROC of 0.934. However, these high values are overestimated and misleading due to the inherent imbalance of the data. Note that the number of epitope-interacting TCRs follows an extreme long-tail distribution (Fig. 3A, 1B, and Supplementary Figure S3) that most TCRs (70%) are associated with the top 5% epitopes. As a result, *Random Epitope* would produce skewed negative data that most majority of epitopes were matched with far more negative TCRs than positive TCRs (Supplementary Figure S3). Trained with such a skewed dataset, TEINet was driven to make predictions based on the epitope sequences without the participation of TCRs, as discussed in Dens *et al*. [32]. That is, when the input pairs consist of frequent epitopes, the model tends to predict “1s”, and conversely, it is likely to predict “0s” when encountering pairs with infrequent epitopes. Thus, TEINet with *Random Epitope* obtains a misleading high performance: (1) for TCRs, TEINet often predicts high scores when they are linked to frequent epitopes, which results in high Precision, Recall, and NDCG since frequent epitopes appear in most paired samples; (2) for epitopes, TEINet tends to predict high scores for pairs with frequent epitopes that possess abundant positive binding TCRs and sparse negative binding TCRs, and low scores for pairs with rare epitopes that are linked to abundant negative TCRs and sparse positive binding TCRs, which leads to high AUROC. Indeed, TEINet with *Random Epitope* obtained high prediction scores for both positive and negative pairs with frequent epitopes (Fig. 3B). For instance, it outputs an average prediction score of 0.99 and 0.93 for respective positive and negative pairs of the most frequent epitope (KLGGALQAK). As a result, those negative pairs will be classified as false positives in the generic performance evaluation. Moreover, due to the long-tail distribution of the epitope-associated TCRs, *Random Epitope* will generate far fewer negative pairs than positive pairs, so that those false positives have minor impact on the generic performance evaluation, resulting in a misleading high AUROC.

**Figure 3:**
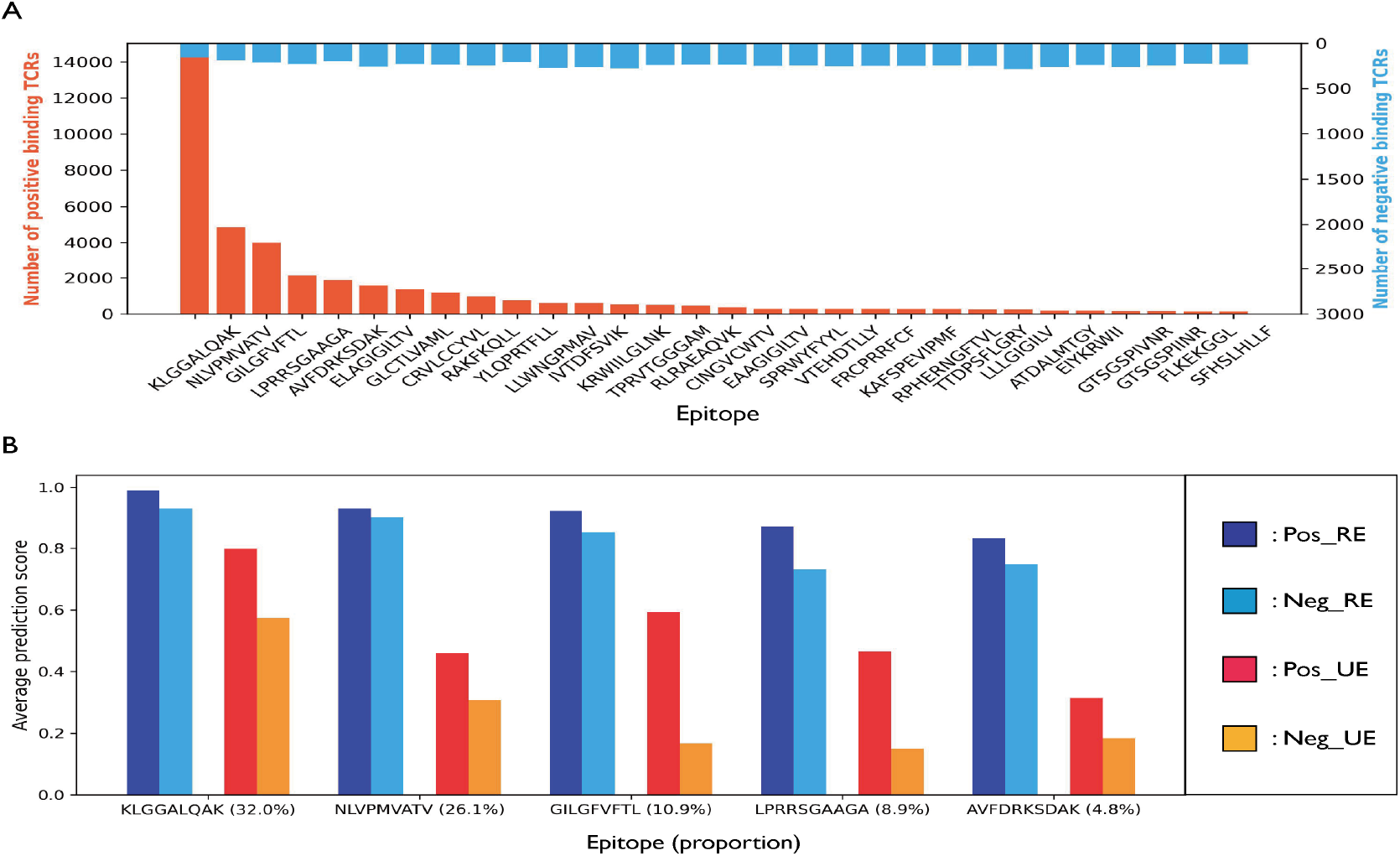
Distribution of the number of epitope-associated TCRs and the average prediction scores for them. (A) Distribution of the number of positive and negative TCRs sampled by *Random Epitope* for the 30 most abundant epitopes. Given that the epitopes are sampled randomly for each TCR and the epitope-associated TCRs follow an extreme long-tail distribution, there are far fewer negative samples than positive samples for abundant epitopes, whereas for most epitopes, there are far more nagative samples than positive samples. (B) The average prediction scores for the positive and negative pairs of the top 5 most abundant epitopes. “Pos” and “Neg” stand for positive and negative samples; “RE” and “UE” represent *Random Epitope* and *Unified Epitope*. We observed that for both positive and negative pairs of abundant epitopes, *Random Epitope* will produce high predictive scores. Such a problem is greatly relieved by *Unified Epitope*.

Overall, our results and analysis indicate that *Unified Epitope* is more appropriate for negative sampling in the TCR-epitope prediction task, which is further supported in the evaluation on independent datasets (see the following section). To eliminate potential model bias introduced by TEINet, we performed the same experiments using the ImRex model and obtained similar results (Supplementary Table S1). In the remaining experiments, *Unified Epitope* is selected as the default strategy.

### Performance of TEINet

To assess the performance of TEINet, we compared it with three existing approaches: ImRex [11], TITAN [12] and pMTNet [14]. ImRex encodes TCRs and epitopes based on their physicochemical properties and utilizes a CNN (convolutional neural network) to process the combined encodings. Similar to our proposed TEINet, TITAN and pMTNet both make use of the pretrained encoders. The pMTNet additionally incorporates the information of the MHC allele associated with the epitope to make the prediction.

Figure 4A and 4B show the AUROC and AUPRC of TEINet as well as the three baseline models. TEINet outperforms the baseline methods with an AUROC of 0.760 and an AUPRC of 0.321, while the second best comparative model ImRex has an AUROC of 0.714 and an AUPRC of 0.269. Moreover, we calculated the Precision, Recall, and NDCG of ImRex and still observed superior performance of TEINet (Supplementary Table S1). With learnable encoders that possess the capability of processing sequences in any length, TEINet can better extract sequence information and consequently make more accurate predictions. To investigate whether the superiority of TEINet retains when the similarity of TCRs between training and evaluation datasets decreases, we filtered out pairs in the test set with specific TCRs according to the Levenstein similarity thresholds (Supplementary Text S4). Figure 4C demonstrates the corresponding performance of each model under different similarity thresholds. Again, TEINet outperforms other baseline models. Further, to resolve the concern that pairs consisting of frequent epitopes would dominate the effects on performance, we report the per-epitope AUROC derived by evaluating on paired data for one specific epitope in Fig. 4D. We found no explicit correlation between the AUROC and the number of training samples, indicating that the complexities of the binding pattern for each epitope are different. Similar results were also found in Moris *et al*. [11]. Besides, TEINet is still superior, with the ImRex lagging behind for most epitopes (Fig. 4D).

**Figure 4:**
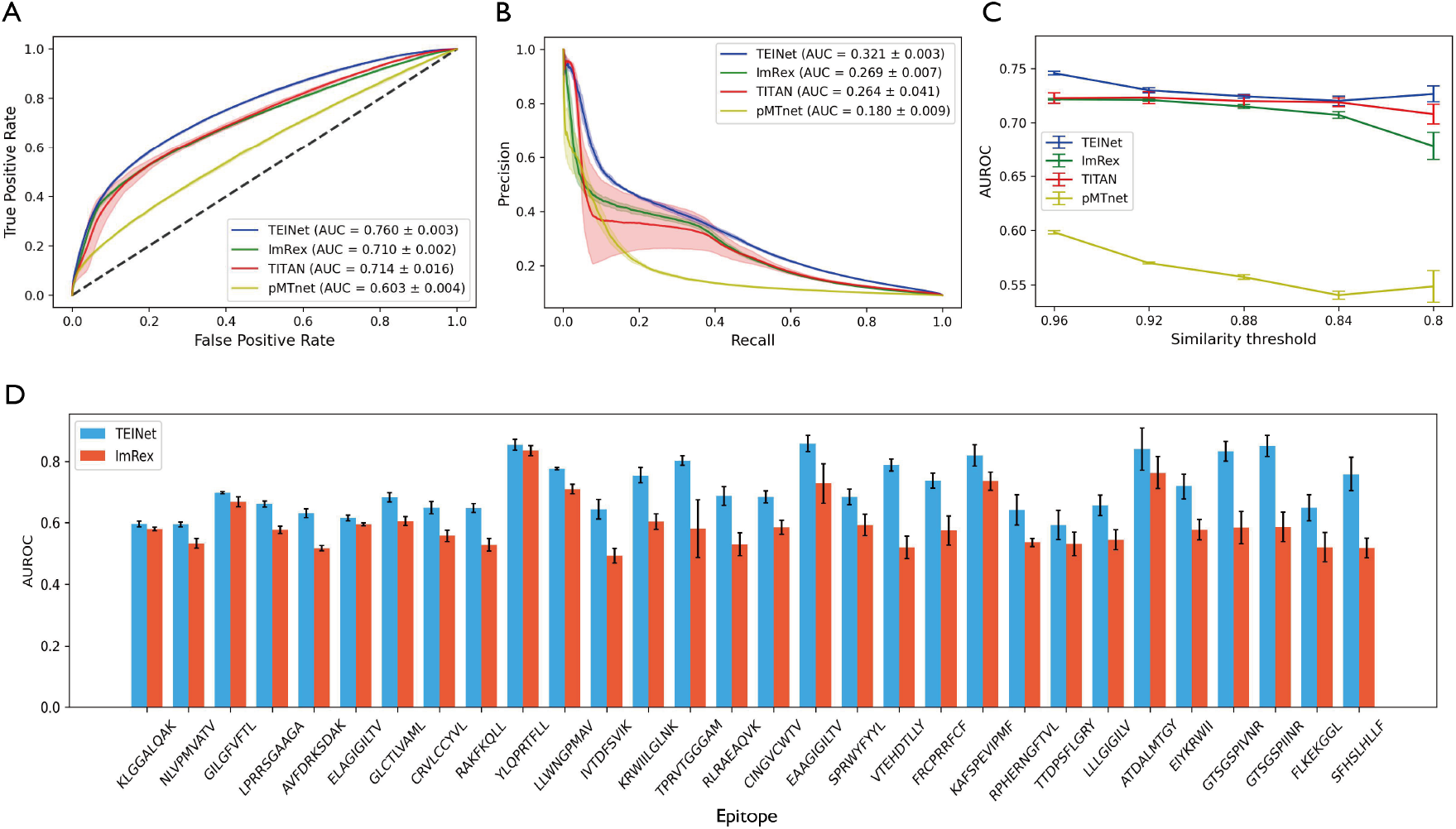
Performance of TEINet and the three baseline models. (A) The receiver operator characteristic (ROC) curves for each model. The area under the ROC curve (AUROC) values are shown in the legend. (B) The precision-recall (PRC) curves for each model. The area under the PRC curve (AUPRC) values are shown in the legend. (C) The AUROC for each model according to different similarity thresholds for filtering the test set. (D) The per-epitope AUROC performance for the top 30 most abundant epitopes. TEINet outperforms ImRex in these epitopes.

### Impact of pretraining

Transfer learning is becoming an integral part of the design of deep learning models for the prediction of TCR binding specificity. Recently developed models tend to employ pretrained encoders to transform amino acid sequences into vector representations [12, 14, 15, 19]. An analysis of the impact of the pretraining step is in demand to provide a better understanding of the pretrained encoders.

First, without the pretraining step, the performance of TEINet dropped significantly with an AUROC of 0.675, which demonstrated the necessity of the pretraining step. Next, we explored the influence of the TCR and epitope encoders singly and simultaneously (Fig. 5A-C). It is clear that the pretraining of TCRs greatly enhanced the model performance (Fig. 5A), whereas the pretraining of epitopes only brought about slight and unstable improvement (Fig. 5B). Given that the diversity of TCRs (41,610 unique samples) is much higher than that of epitopes (180 unique samples), pretraining of TCRs enables them to be distributed separably in the feature space, which is more important for making a prediction. Further, these two encoders improved the performance synergistically and achieved the best performance (Fig. 5C). Utilizing both pretrained encoders enhanced the AUROC by around 0.01 than using the TCR encoder alone. Notably, we observed that when the pretraining of the TCR encoder exceeded a certain epoch, the final performance dropped (Fig. 5A and C). Thus, the degree of the pretraining needs to be tuned carefully; otherwise, the model might have the problem of overfitting.

**Figure 5:**
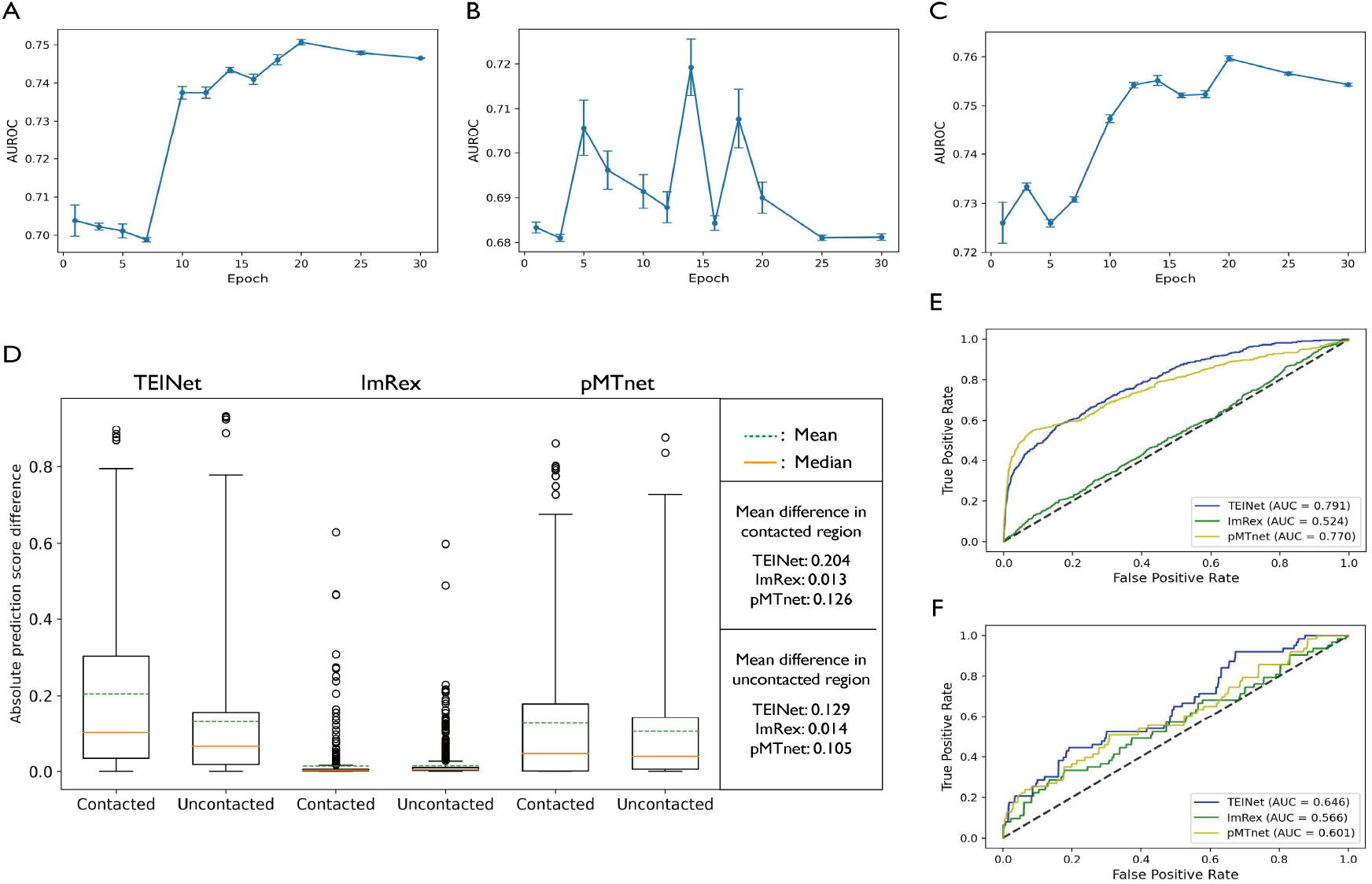
Investigation of the impact of the pretraining stage and further validations of TEINet. (A-C) The AUROC values with the encoders trained for different epochs. (A) Only pretrain the encoder for TCRs. (B) Only pretrain the encoder for epitopes. (C) Pretrain the encoders for TCRs and epitopes. (D) The absolute difference of prediction scores for each model between the contacted and uncontacted residues. Using TEINet, residues with direct contacts are more likely to induce larger changes in the predicted binding strength than non-contact residues. (E and F) The ROC curves for each model in the two independent test sets: (E) TBAdb and (F) PDB. The corresponding AUROC values are shown in the legend.

### Structural analysis

Perturbation (mutational) analysis can be used to detect the important amino acid residues for the model prediction [14, 24, 33]. We grouped TCR residues by whether or not they formed any direct contact with any residue of epitopes within 5*Å* and assumed that substitutions inside the contact region would lead to dramatic changes in the predicted binding score. To analyze the effects of predictive models on the contact/non-contact region, we collected 105 solved TCR-epitope interacting complex structures from the public RCSB Protein Data Bank (PDB) database [34] as the ground truth data. We performed the alanine scanning technique in biophysics studies [35] on the TCRs in the PDB database using the predictive models. Figure 5D illustrates the average score difference for each model inside the contact and non-contact region. We observed that for TEINet, the contact residues were more likely to induce larger drops in predicted TCR-epitope binding strength than non-contact residues, which supports our assumption.

### Evaluation on independent datasets

To further compare the predictive performance of each model, we collected two independent test sets. We selected the TBAdb [36] dataset, which includes 439 binding pairs on 414 unique TCRs and 42 epitopes as our first independent test set; The 105 interacting pairs extracted from the PDB database aforementioned were selected as the second independent test set. As before, the same filtering procedure was applied to them. Figure 5E and 5F show the performance of each model on the independent test sets. Again, TEINet achieved superior performance over the other baseline methods. Note that for the PDB dataset, TEINet obtained a lower AUROC value of 0.646. We attributed it to the small overlap of epitopes as there is only 1 epitope in the PDB dataset that also appears in the training data. Moreover, given that the PDB dataset is an approximately balanced dataset with each epitope binding with 1 or 2 TCRs, the *Random Epitope* and *Unified Epitope* will generate similar negative data, which enables us to compare these two strategies by the AUROC value. Thus, we trained two TEINets each using *Random Epitope* or *Unified Epitope* during the training process and then evaluated them on the PDB database constructed with *Random Epitope*. We observed that TEINet trained with *Random Epitope* obtained an AUROC of 0.572, which was surpassed by *Unified Epitope* by a large margin with an AUROC of 0.644 (Supplementary Figure S4). This finding further supports the advantage of *Unified Epitope*.

## 1 Discussion

The prediction of TCR specificity to epitope has been a challenging problem. The immense searching space of immune receptors, lack of curated training samples, and absence of negative samples remain issues for algorithm development. In recent years, public databases have been accumulating an enormous amount of TCR-epitope interacting data. Benefiting from the enrichment of data enrichment of available data, it is possible to develop accurate deep learning models to tackle the challenging task of TCR-epitope interaction prediction.

In this work, we have proposed TEINet, a new deep learning model for predicting the TCR binding specificity. TEINet only requires the CDR3*β* chain of the TCR and epitope sequence of the pMHC complex to make the prediction. Though the CDR3*α* chain and the MHC allele are shown to be beneficial in this task [10, 13–15, 23], the paired data is still rare compared to single-chain data, which limits the generalizability of the pair-chain model. We leave the exploration of both CDR3 chains and MHC alleles in future work. TEINet employed the TCRpeg [24], a deep autoregressive model, to extract the sequence information of TCRs and epitopes and transform them into numerical vector space. The TCRpeg was pretrained in a self-supervised learning manner on large-scale sequence data to learn a more general pattern to encode TCRs/epitopes. TEINet then combined the encodings of TCRs and epitopes and used a fully-connected neural network to make the final prediction, leveraging the knowledge from TCRs and epitopes.

To train and evaluate a supervised model, negative samples are required. However, currently there is no unified method for negative sampling, which poses a challenge for comparing different models. For example, *Random TCR* was applied in pMTNet [14]; *Reference TCR* was applied in NetTCR [15, 18]; *Random Epitope* was employed in TITAN [12]; *Unified Epitope* was employed in ImRex [11]. We thus proposed three metrics, Precision, Recall, and NDCG that are unrelated to negative samples to compare different sampling strategies. We manifested that *Unified Epitope* is the winner among these four sampling schemas for the development of a more accurate model, given that it achieved superior Precision, Recall, and NDCG among the first three schemas and that *Random Epitope* breaks the uniformity between positive and negative data, which leads to misleading performance. Thus, we recommend *Unified Epitope* as the default negative sampling method in future works.

To showcase the predictive strength of TEINet, we compared TEINet with another three published deep learning models: ImRex [11], TITAN [12], and pMTNet [14]. We performed the 5-fold cross-validation procedure on our constructed dataset which consists of 44,682 interacting pairs. We observed that TEINet achieved an AUROC of 0.760 and an AUPRC of 0.321 and outperformed other comparative models with the best AUROC of 0.714 and AUPRC of 0.269. Further, we also evaluated and compared these models on two additional independent test sets. Again, TEINet surpassed other baseline models.

The usage of the transfer learning technique has become a trend in the design of deep learning models for the TCR-epitope binding prediction task. Instead of using the physicochemical properties of amino acid sequences to construct the features of TCRs and epitopes, many recently published models capitalized on the pretrained encoders that leveraged the knowledge learned from other tasks with abundant data [12, 14, 15, 19]. However, the impact of the pretraining step on the final prediction accuracy remains unknown, which could potentially hinder the exploitability of pretrained encoders. Here, we disentangled the effect from each encoder (Fig. 5A-5C). We first observed that the pretraining of the TCR encoder improved the TEINet by a much larger margin than that of the epitope encoder, which could be explained by the vast diversity of TCRs. More importantly, we found that excessive pretraining might harm the performance, so that the degree of pretraining needs to be tuned carefully.

At last, we analyzed whether the prediction from TEINet can reveal the structural information of the interacting complex. We grouped residues of TCRs that form any contact with epitope within 5*Å* into the contact region. Contact residues should be more important than non-contact residues in forming the interaction between TCRs and epitopes [37]. Indeed, larger drops of predicted scores were observed inside the contact region than non-contact region using TEINet.

In summary, we have designed TEINet to predict the interaction between TCRs and their epitope targets. Our results demonstrate that TEINet achieved superior performance over three other comparative models only by using the information of CDR3*β* chains and epitope sequences. We also compared different negative sampling strategies and suggested that *Unified Epitope* is more appropriate for the development of a generalized model. We expected that with enhanced accuracy in predicting the potential immune response of T-cells to epitopes, TEINet could be beneficial for the *in silico* design and implementation of immunotherapy in the era of personalized medicine.

## Acknowledgments

We thank all contributors to VDJdb, McPAS-TCR, and other TCR specificity datasets for making their data publicly available.

## Notes

### Competing Interest Statement

The authors have declared no competing interest.

