## Supplementary Materials for "TEINet: a deep learning framework for prediction of TCR-epitope binding specificity"

Yuepeng Jiang[1], Miaozhe Huo[1] and Shuai Cheng Li[1*]
[1]Department of Computer Science, City University of Hong Kong, Kowloon Tong, Hong Kong

#### 1 Text S1

The Discounted Cumulative Gain (DCG) takes the ranking of the top k predictions into account, which penalizes the situations that the prediction scores for the true binding pairs are located at the later part among the top k binding predictions. The formula of DCG accumulated at a particular rank position k is defined as:

$$DCG@k = \sum_{i=1}^{p} \frac{rel_i}{log_2(i+1)}, \tag{1}$$

where $rel_i$ is the graded relevance of the result at position $i$. Since the prediction task is binary (either binding or non-binding), the graded relevance is also binary: $rel_i \in \{0, 1\}$. The ideal DCG (IDCG) is the the maximum possible DCG through position k, which can be formulated as:

$$IDCG@k = \sum_{i=1}^{T} \frac{rel_i}{log_2(i+1)}, \tag{2}$$

where T is the number of the true binding epitopes for a given TCR. Then the normalized discounted cumulative gain (NDCG) is computed by:

$$NDCG@k = \frac{DCG@k}{IDCG@k}. \tag{3}$$

NDCG is more stable and instructive for evaluating the average performance of the predictive model in discovering the binding epitopes for TCRs and is commonly used in Recommender System.

### 2 Text S2

TCRpeg [1] is an autoregressive model that formulates the probability of a TCR sequence $\boldsymbol{x}$ as $p(\boldsymbol{x}|\boldsymbol{\theta})$, where the parameters $\boldsymbol{\theta}$ capture the latent evolutionary patterns to generate $\boldsymbol{x}$. The probability density $p(\boldsymbol{x}|\boldsymbol{\theta})$ can be calculated by the product of probabilities conditioned on previous residues along a sequence with length $L$ through an autoregressive likelihood

$$p(\boldsymbol{x}|\boldsymbol{\theta}) = p(x_1|\boldsymbol{\theta}) \prod_{i=2}^{L} p(x_i|x_1, ..., x_{i-1}; \boldsymbol{\theta}). \tag{4}$$

TCRpeg utilized gated recurrent units (GRUs), to model the autoregressive likelihood and after the training process, the hidden layer of TCRpeg can be extracted as the numerial embeddings for input TCR (epitope) sequences. To see the details of TCRpeg, please refer to the original paper.

### 3 Text S3

**Training of Encoders.** We first trained the word2vec algorithm on the TCR and epitope sequences separately and obtained two sets of embeddings for amino acids in different domains (TCR vs. epitope). Specifically, for TCRs we trained for 10 epochs with a batch size of 64 and learning rate of 0.0001. The window size and the embedding size are set to 2 and 64, respectively. For epitopes, we trained word2vec under the same settings used in TCRs. The TCRpeg-TCR and TCRpeg-Epi both have 3 hidden layers each with 768 neurons. They were trained for 20 epochs with a learning rate of 0.001 and a batch size of 32.

### 4 Text S4

To avoid remembering the training set for prediction and perform a more robust model evaluation, we filter the test set based on the Levenshtein similarity score: for a pair $(t_i, e_i)$ in the test set, if $t_i$ possess a levenshtein similaity score $L_sim(t_i, t_i^{'})$ with any TCR $t_i^{'}$ in the training set, then this pair will be filtered out. The levenshtein similarity score is defined as:

$$L_sim(t_1, t_2) = 1 - \frac{Dis(t_1, t_2)}{len(t_1) + len(t_2)}, \tag{5}$$

where $Dis(t_1, t_2)$ computes the Levenshtein distance (edit distance) between sequences $t_1$ and $t_2$.

### 5 Supplementary Table

Table S1: The Precision, Recall, and NDCG of each negative sampling method using ImRex model. Performance is shown with the best performance among *Reference TCR*, *Random TCR*, and *Unified Epitope* in bold.
Compared to TEINet, ImRex shows significantly lower performance.

| Method | Precision@3 | Recall@3 | NDCG@3 | Precision@10 | Recall@10 | NDCG@10 |
|---|---|---|---|---|---|---|
| *Reference TCR* | 0.003±0.000 | 0.009±0.001 | 0.006±0.001 | 0.005±0.000 | 0.046±0.001 | 0.0194±0.002 |
| *Random TCR* | 0.002±0.000 | 0.006±0.000 | 0.005±0.004 | 0.003±0.001 | 0.024±0.002 | 0.012±0.002 |
| *Unified Epitope* | **0.051±0.001** | **0.151±0.003** | **0.118±0.004** | **0.032±0.004** | **0.315±0.011** | **0.176±0.009** |
| *Random Epitope* | 0.177±0.003 | 0.519±0.006 | 0.433±0.005 | 0.074±0.002 | 0.720±0.003 | 0.505±0.005 |

### 6 Supplementary Figures

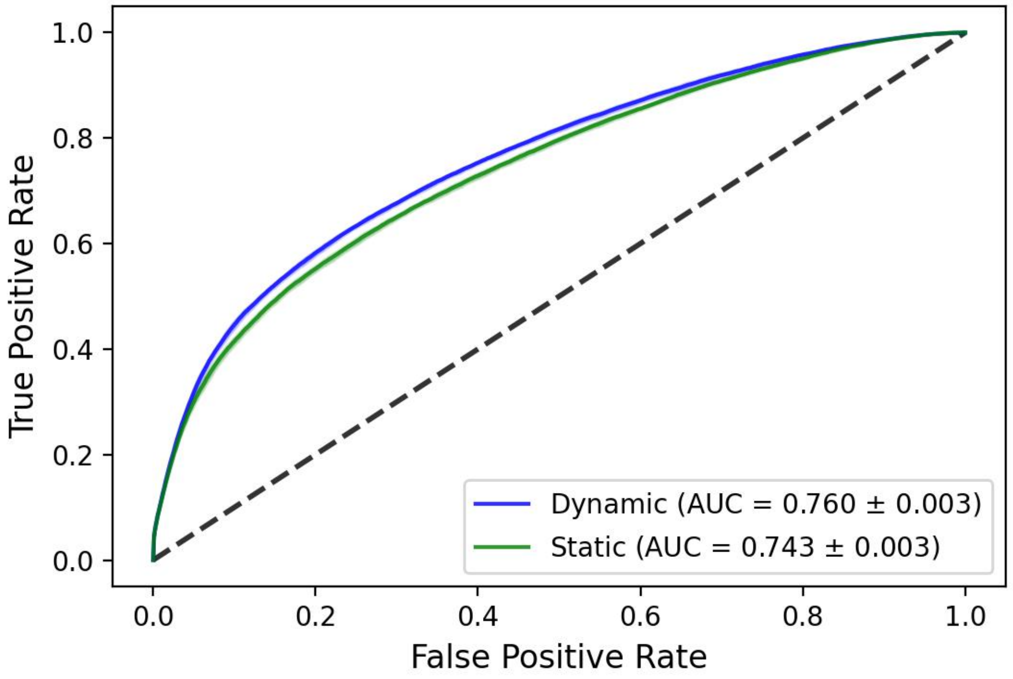


Figure S1: The average performance of dynamic sampling strategy and static training strategy. We found that sampling the negative pairs during the training process can improve the model performance.

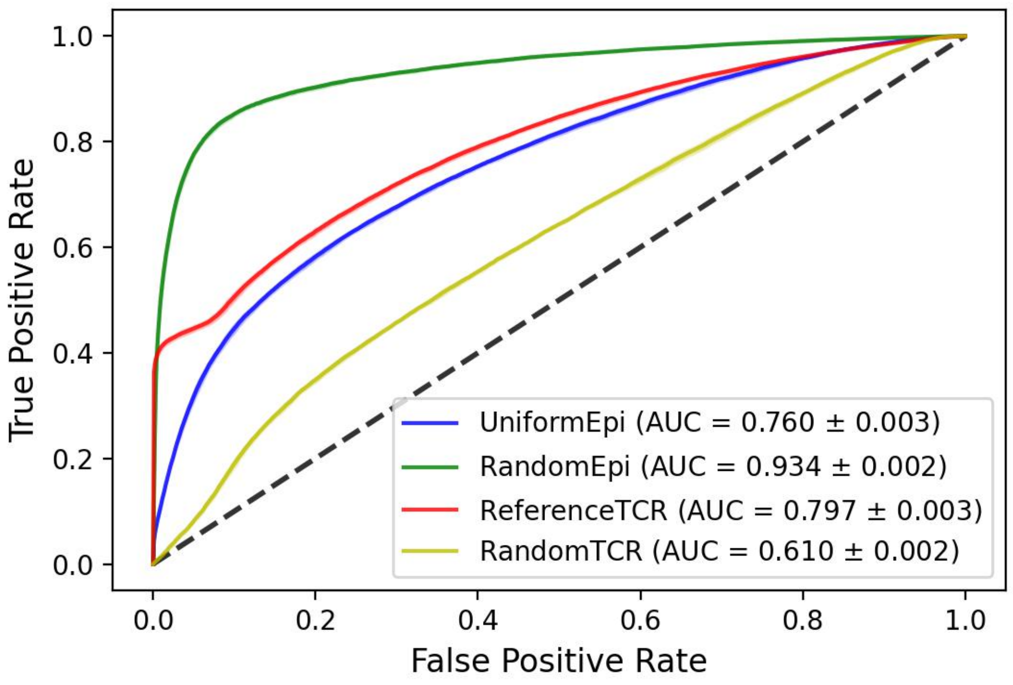


Figure S2: The average AUROC performance for the four negative sampling strategies. We can see that different strategy will have AUC value in different scales. Thus, a unified strategy is needed for model comparison.

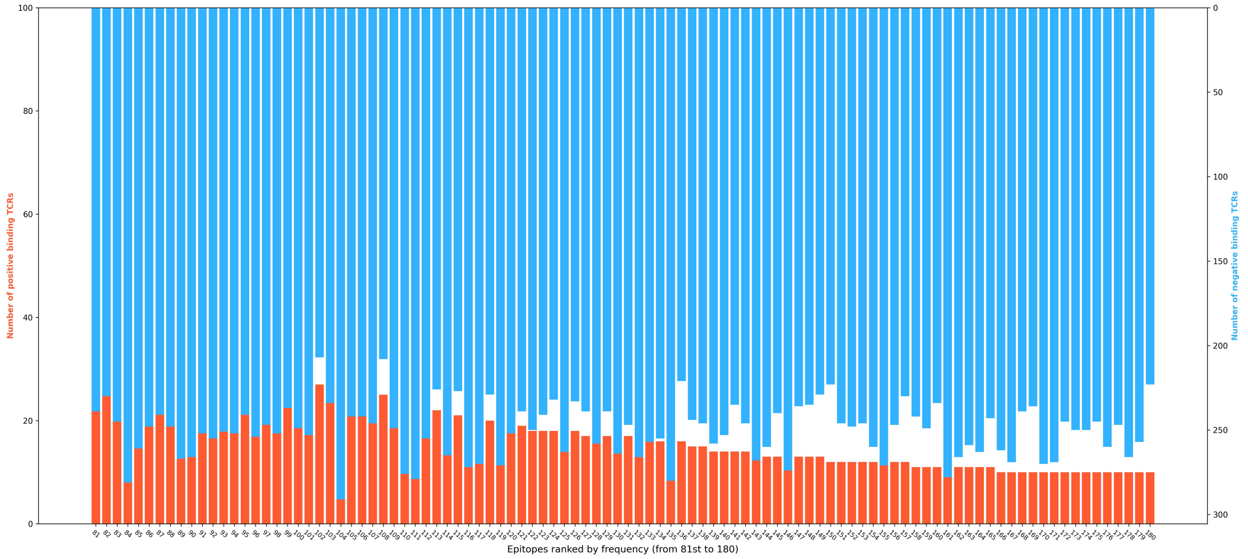


Figure S3: Distribution of the number of positive and negative TCRs sampled by *Random Epitope* for the last 100 abundant epitopes (from 81st to 180th). Given that the epitope-associated TCRs follow an extreme long-tail distribution, there are far more negative samples than positive samples for these epitopes.

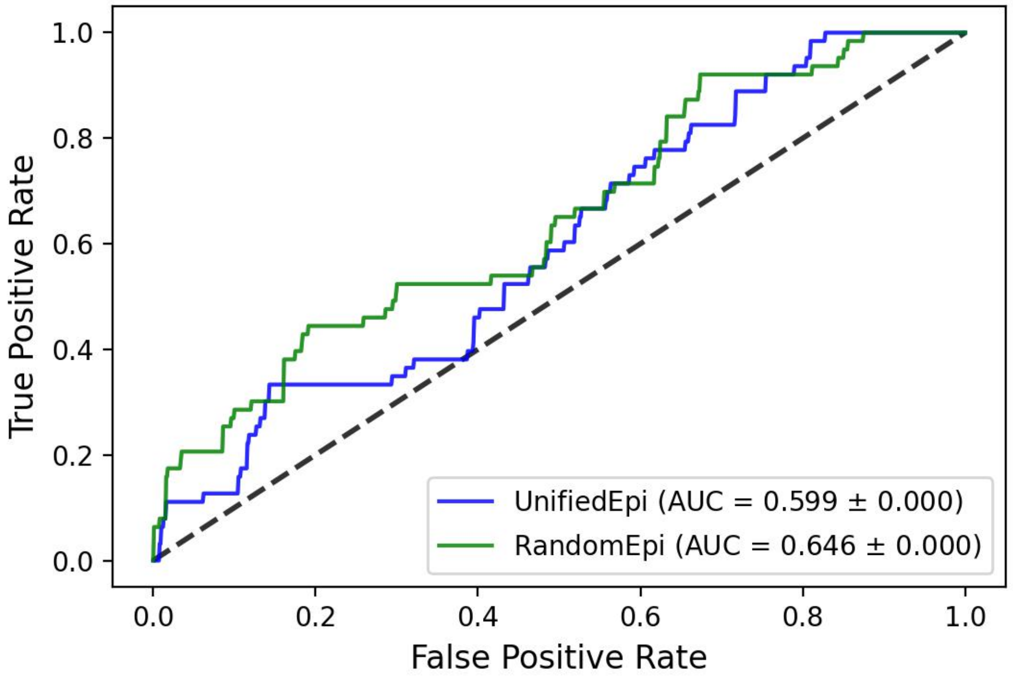


Figure S4: The performance of *Unified Epitope* and *Random Epitope* on the PDB database using TEINet. Since the PDB database is a nearly balanced dataset, these two sampling strategies will produce similar negative samples. Thus, we can now use the AUC value to compare these two strategies.

#### References

[1] Yuepeng Jiang and Shuai Cheng Li. Deep autoregressive generative models capture the intrinsics embedded in t-cell receptor repertoires. *bioRxiv*, 2022.
